# The architecture of the human RNA-binding protein regulatory network

**DOI:** 10.1101/041426

**Authors:** Alessandro Quattrone, Erik Dassi

## Abstract

RNA-binding proteins (RBPs) are key players of post-transcriptional regulation of gene expression. These proteins influence both cellular physiology and pathology by regulating processes ranging from splicing and polyadenylation to mRNA localization, stability, and translation. To fine-tune the outcome of their regulatory action, RBPs rely on an intricate web of competitive and cooperative interactions. Several studies have described individual interactions of RBPs with RBP mRNAs, suggestive of a RBP-RBP regulatory structure. Here we present the first systematic investigation of this structure, based on a network including almost fifty thousand experimentally determined interactions between RBPs and bound RBP mRNAs.

Our analysis identified two features defining the structure of the RBP-RBP regulatory network. What we call “RBP clusters” are groups of densely interconnected RBPs which co-regulate their targets, suggesting a tight control of cooperative and competitive behaviors. “RBP chains”, instead, are hierarchical structures driven by evolutionarily ancient RBPs, which connect the RBP clusters and could in this way provide the flexibility to coordinate the tuning of a broad set of biological processes.

The combination of these two features suggests that RBP chains may use the modulation of their RBP targets to coordinately control the different cell programs controlled by the RBP clusters. Under this *island-hopping* model, the regulatory signal flowing through the chains hops from one RBP cluster to another, implementing elaborate regulatory plans to impact cellular phenotypes. This work thus establishes RBP-RBP interactions as a backbone driving post-transcriptional regulation of gene expression to allow the fine-grained control of RBPs and their targets.

## Introduction

In the last years post-transcriptional regulation of gene expression (PTR) has gained recognition as a crucial determinant of protein levels, and consequent cell phenotypes (Schwanhäusser et al. 2011; Vogel et al. 2010), stimulating a rising interest in studies focused on RNA-binding proteins (RBPs) and the interactions with their RNA targets.

RBPs are a key class of regulators in PTR. They are less than two thousand proteins in the human genome (almost 1200 verified RBPs plus several recently discovered ones (Castello et al. 2012)) and are made of modular domains of which RRM is the most represented one, found in over 200 proteins (Lunde et al. 2007). RBPs control processes ranging from splicing and polyadenylation to mRNA localization, stability, and translation (Gerstberger et al. 2014). To fine-tune the outcome of their regulatory action, RBPs rely on an intricate web of competitive and cooperative interactions (Dassi 2017).

Techniques such as ribonucleoprotein immunoprecipitation (RIP) and cross-linking and immunoprecipitation (CLIP) variants (Lee and Ule 2018) now allow us to identify the RNA targets of an RBP at the genome-wide scale. RBPs are involved in multiple aspects of physiology (e.g. brain and ovary development, immune response and the circadian cycle (Gerstberger et al. 2014; Lim and Allada 2013)) and pathology, being their alteration associated with a variety of diseases such as cancer, neurological and neuromuscular disorders (Wurth and Gebauer 2015; Lukong et al. 2008). The importance of obtaining a proper understanding of RBP properties and functions is thus evident.

While identifying the mRNA targets of RBPs, several works have highlighted among them an enrichment of mRNAs coding for gene expression regulators, including other RBPs but also transcription factors (TFs). This finding brought to the *regulator-of-regulators* concept (Keene 2007; Mansfield and Keene 2009), hinting at the existence of an extensive regulatory hierarchy of RBPs. For instance, we and others have specifically studied the *HuR*/*ELAVL1* protein (Dassi et al. 2013; Mukherjee et al. 2011; Pullmann et al. 2007), which resulted to regulate the mRNAs of many RBPs (Mukherjee et al. 2011), several of which contain its same RNA-binding domain, the RRM. The increasing number of high-throughput data available is now allowing us to probe if this phenomenon occurs on a genome-wide scale. We chose to address this issue by specifically extracting the binding map of RBPs to their cognate mRNA and mRNAs of other RBPs. A similar approach has been previously applied for TF targets and metabolic networks in lower organisms such as *E. coli* and *S. cerevisiae* (Qijun Liu et al. 2009; Jothi et al. 2009; Pham et al. 2007); the human TF-TF regulatory interaction network, testing the *regulator-of-regulators* concept in TFs, has also been described for 41 cell types (Neph et al. 2012).

We present here the first systematic characterization of the RBP regulatory network, built by integrating experimental data on RBP targets. While sharing several properties of gene regulatory networks, its distinctive local structure hints at the specific dynamics of post-transcriptional regulation. We identified two major components which define the network structure. First, we found groups of densely connected RBPs which control each other to likely regulate cooperative and competitive behaviors on mutual targets. Then, we identified hierarchical node chains as the second feature shaping the network. In combination with RBP groups, these widespread regulatory units concur to the formation of a post-transcriptional backbone acting on multiple processes at once and could serve to coordinate major cell programs shaping cell phenotypes.

## Results

### Building the RBP-RBP network

Large-scale mapping of interactions between RBPs and their cognate mRNAs has been conducted by CLIP-like approaches (Lee and Ule 2018) in a few cellular systems, primarily HEK293, HeLa, and MCF7 cell lines. We previously collected these and other interactions in the AURA 2 database (Dassi et al. 2014). We have now built the human RBP-RBP mRNA interaction network by extracting all related data and filtering each interaction by the expression of both interactors in the HEK293 cell line. To verify the generality of properties identified in this cell line, we have also constructed the same network for the other two cell lines with sufficient CLIP-like data, HeLa and MCF7. In our network vertices represent RBPs, and the presence of an edge between a source (protein) and a target (mRNA) RBP implies binding of the target RBP mRNA by the source RBP (which could result in post-transcriptional control of gene expression). The network includes 1536 RBPs out of 1827 (see methods for details on how we built the RBP list) connected by 47957 interactions. A total of 176 RBPs (11,5%) have outgoing interactions in the network (i.e., they bind the mRNA of an RBP) mostly coming from CLIP-like assays; the median network degree (number of connections) is 29, while the median number of individual binding sites for each RBP on each target RBP mRNAs is equal to 4. Among RBPs with outgoing interactions, 63 (35.8%) have self-loops (i.e., they bind their mRNA), confirming the general propensity of RBPs for autologous regulation. All interactions are listed in **Table S1**, and an interactive browser allowing to explore this and other networks is available at the AURA 2 (Dassi et al. 2014) website (http://aura.science.unitn.it).

### The RBP-RBP network is a navigable “small-world” network

We first sought to verify whether the RBP-RBP network is a typical gene regulatory network, i.e. is “scale-free” and “small-world”. To this end, we computed several global properties of the HEK293 cell network, shown in **Figure 1**. The degree distribution (**1A**) is following a power-law, with most nodes having a degree lower than 50 and a minor fraction reaching degrees over 200. This suggests that the network is scale-free, composed of a few central hubs and many progressively more peripheral nodes. The diameter (**1B,** D=5) indicates the network to be largely explorable by a few steps. Clustering coefficients (**1B**) suggest the presence of local-scale clustering (1-neighbor coefficient, CC1=0.507) which is lost when extending to more distant nodes (2-neighbor coefficient, CC2=0.0115). Eventually, closeness centrality (**1B**, C_c_=0.5033) reiterates that most nodes are reachable by a small number of steps. We thus quantified this intuitive idea of network small-worldness by computing the *S^WS^* measure (Humphries and Gurney 2008), which classifies a network as small-world when greater than 1. We obtained a value of 31.03, clearly supporting the hypothesis. Taken together, these values indeed put the network into the “small-world” class. Given its small diameter and high connectedness, the network can be considered navigable (Kleinberg 2000), i.e. apt to promote efficient information transmission along its paths. Eventually, we investigated the network control structure (how it can be driven to any of its possible states), as described in (Ruths and Ruths 2014). We computed the network control profile, which resulted being [s=0.00367, e=0.99632, i=0.0], with *s* representing sources, *e* the external dilations and *i* the internal dilations. Hence, the network is dominated by external dilations (*e*), a fact that locates it in the class of top-down organization systems, structured to produce a correlated behavior throughout the system: members of this class are transcriptional networks, peer-to-peer systems, and corporate organizations (Ruths and Ruths 2014). These properties also hold in the HeLa and MCF7 networks, suggesting the stability of the network structure with different subsets of expressed RBPs (**Table S2**). We thus focused on the HEK293 network only for subsequent analyses.

**Figure 1:**
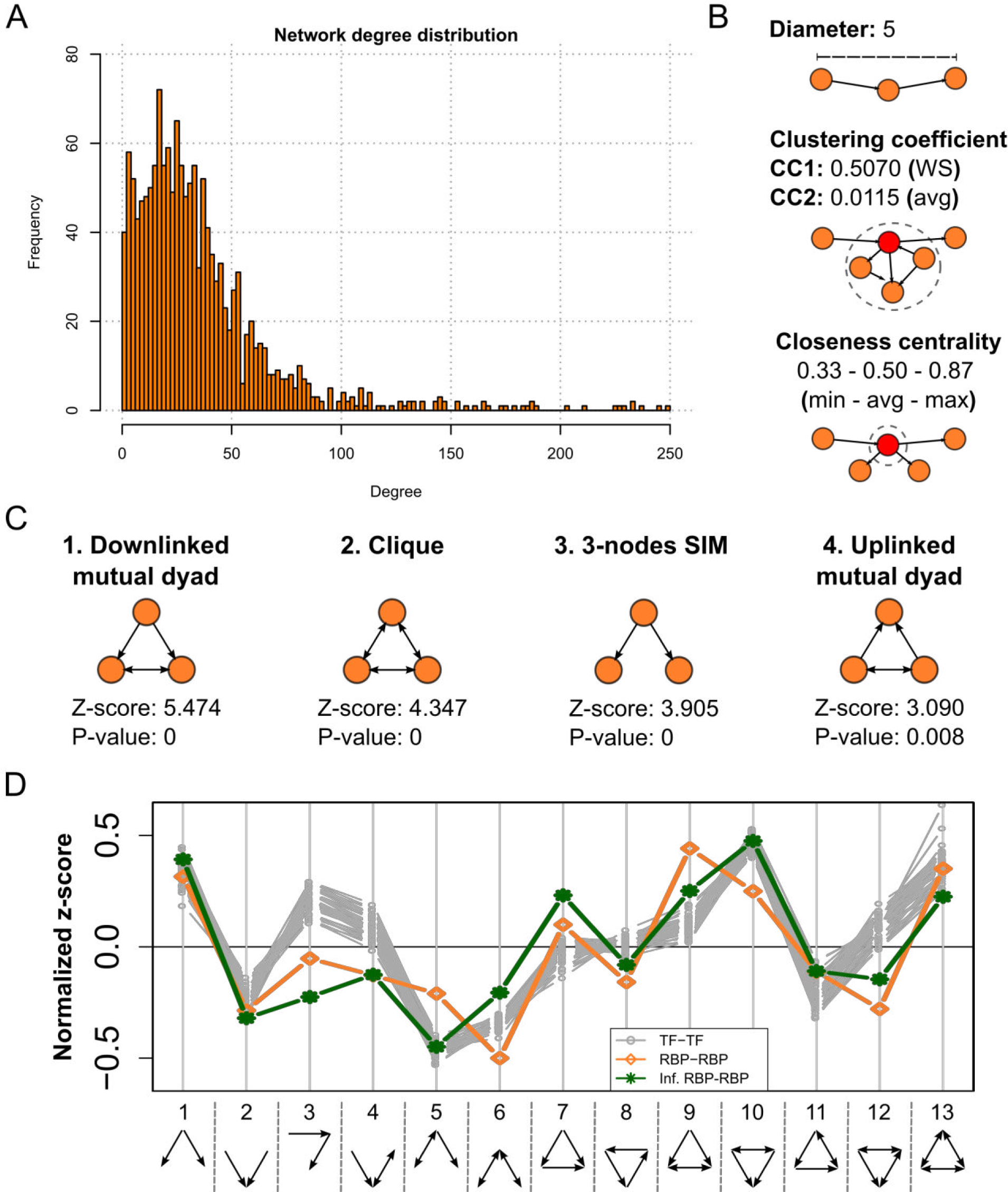
The RBP-RBP network is a gene regulatory network with a distinctive structure. **A)** shows the network degree distribution (up to 250), which follows a power-law. **B)** shows the network diameter (top), its average clustering coefficients (middle, Watts-Strogatz 1-neighbor coefficient, named CC1, and 2-neighbor coefficient, named 115 CC2) and closeness centrality (bottom, minimum, average and maximum values for all nodes). **C)** shows the four most significant 3-nodes motifs identified by FANMOD with their z-score and p-value. **D)** displays the triad significance profile for the RBP-RBP network (orange line), the inferred RBP-RBP network (green line) and 41 TF-TF networks (gray lines). Positive z-scores indicate enrichment, negative ones depletion. While most motifs have similar z-scores in both networks, motifs 3, 4, 5, 9, 10 and 12 are differentially enriched in the RBP-RBP network, suggesting a distinctive structure with respect to the TF-TF networks.

### RBP-RBP interactions define a hierarchical network structure

We then analyzed the local network structure by identifying motifs, i.e. recurrent patterns of RBP interaction. We used FANMOD (Wernicke and Rasche 2006) to look for 3-nodes motifs, of which several patterns have previously been characterized (e.g., the feed-forward loop and others (Milo et al. 2002)). The most significant motifs are shown in **Figure 1C**: among these, the *down-linked mutual dyad* (DMD) is the most enriched motif in our network. Together with the *single-input module* (SIM, third most enriched motif), these motifs indicate widespread use of hub-like patterns. The enrichment in DMD and *up-linked mutual dyad* (UMD, fourth most enriched motif), suggest a structure of ranked clusters for our network. Under this model, the dyads connect different hierarchical ranks within a network, with individual ranks structured as node clusters (1985; de Nooy et al. 2005). Instances of these motifs include *FXR2*, *HNRNPF*, and *TNRC6B* for the DMD, and *IGF2BP1*, *YWHAE,* and *YWHAG* for the SIM (with the first binding to the other two mRNAs). One realization of the UMD is that of *ELAVL1*, *LIN28B*, and *SYNCRIP* (e.g., both binding to *SYNCRIP* mRNA and the mRNA of each other).

We then identified 4-node motifs, the most significant of which are shown in **Supplementary Figure S1**. Among these, the forwarded uplinked mutual dyad is used to forward the output of an uplinked mutual dyad to a further RBP, and thus is a hierarchical, rank-connecting extension of the UMD. Furthermore, the chain-feeding dyad is made of a dyad which transmits its regulatory signal to two linearly connected RBPs, thus realizing a hierarchical structure as well. Given their properties, these two motifs provide further support to a ranked clusters model for the structure of our network.

### The structure of the RBP-RBP network is different from the TF-TF one

We thus sought to compare the motif structure of the RBP-RBP network with the one of another network of regulators, the TF-TF network described in (Neph et al. 2012) for 41 cell types. We thus computed the triad significance profile (TSP) for these networks as described in (Milo et al. 2002). The TSP quantifies the use of the various three-nodes motifs by the network under analysis, and thus recapitulates its local structure. To complement this analysis, we also asked ourselves whether the structure of our network could be considered representative of the unavailable “complete” RBP-RBP network. To answer this question we thus built an inferred RBP-RBP network by collecting experimentally determined RBP-bound regions as per a protein occupancy profiling assay in HEK293 cells (Baltz et al. 2012). We then matched these regions to the binding motifs of 193 human RBPs derived from the *in vitro* RNAcompete assay (Ray et al. 2013). We obtained a network of 108161 RBP-RBP interactions. This network, independently reconstructed from two experimental datasets, becomes a validation of the general structure we propose for the RBP-RBP network.

We eventually compared the TSP of the three networks. The results are shown in **Figure 1D**, and we observe two salient aspects. First, the RBP-RBP network and its inferred version have a very similar motif structure (Pearson correlation=0.838, p-value=3.47e-04), with limited magnitude differences only, suggesting that our network structure is reproducible and a representative cross-section of the complete set of RBP-RBP interactions. Then, the TF-TF structure is instead more distant (mean Pearson correlation=0.719 across the 41 networks). Indeed, 5/13 motifs are differentially represented in the RBP-RBP network (enriched instead of depleted or vice versa), and the DMD is preferred over the UMD (the opposite being true for the TF-TF networks). This suggests specialization of network structures by RBP-RBP interactions with respect to TF-TF ones.

### The stoichiometry of RBP complexes is not determined by RBP-RBP regulatory interactions

We then asked if some type of biological constraints could be behind the evolutionary shaping of the specific geometry of the RBP-RBP network. One likely hypothesis is that many constraints are produced by interacting RBPs being part of the same ribonucleoprotein complex. To test this hypothesis, we overlapped the interactions in our network with the experimental binary protein-protein interactions (PPIs) contained in STRING (Szklarczyk et al. 2017), IntAct (Orchard et al. 2014), and BioPlex (Huttlin et al. 2017). The low amount of network interactions found to be mirrored in PPIs (3.37% for STRING, 0.57% for IntAct, and 0.29% for BioPlex) suggests that the network wiring is not made to assure the availability of RBPs for complex assembly. As this analysis dealt with single interactions, we then turned to whole complexes, as stoichiometric ones (i.e., requiring precise quantities of each of the components for proper functioning) may instead rely on this mechanism. We employed data from CORUM (Ruepp et al. 2010) and found 1818 interactions overlapping a complex, corresponding to only 3.79% of the network. **Table S3** lists complexes with at least five interactions in the network involving their subunits. A few complexes are highly represented, including the large Drosha complex (95% of its subunits are in the network, connected by 88 interactions) and the spliceosome (83% of its subunits and 732 interactions). This suggests that only for some notable exceptions stoichiometry of protein complexes is possibly driving the establishment of interactions in the RBP-RBP network.

### Communities do not globally define the structure of the RBP-RBP network

To obtain a more general understanding of RBP-RBP interactions, we thus asked ourselves whether the network had a modular structure, made of RBP communities aimed at regulating specific biological processes. We used SurpriseMe (Aldecoa and Marin 2013), a tool for the investigation of community structures implementing several algorithms. SurpriseMe is based on Surprise (S) maximization (Aldecoa and Marin 2013), which has been shown to outperform the classic Girvan-Newman modularity measure Q (Newman and Girvan 2004). We used the communities identified by the two best-scoring algorithms implemented in the tool, namely CPM (Palla et al. 2005) and RNSC (King et al. 2004) (S=13698 and 13353 resp.). These algorithms detected a poor degree of modularity in the network: as shown in **Figure 2A**, 89% of the communities are singletons (i.e., formed by a single RBP) and only 8 contain more than 20 RBPs (1 with CPM and 7 with RNSC). Furthermore, both algorithms identified a huge community comprising a substantial portion of the network, suggesting a limited presence of true clustered structures. In that respect, the TF-TF networks appear to be much more modular, with much fewer communities identified as singletons (avg. of 53 vs. 657 for the RBP-RBP network) and higher average community size (avg. of 5 vs. 2.08). Eventually, we explored the enrichment of biological functions in the communities but detected no clear association involving most members of any of these. CPM and SCluster-derived communities are listed in **Table S4A** and **S4B**. Globally, these results suggest that the conventional community definition does not fit well the RBP-RBP network, which may thus be structured differently.

**Figure 2:**
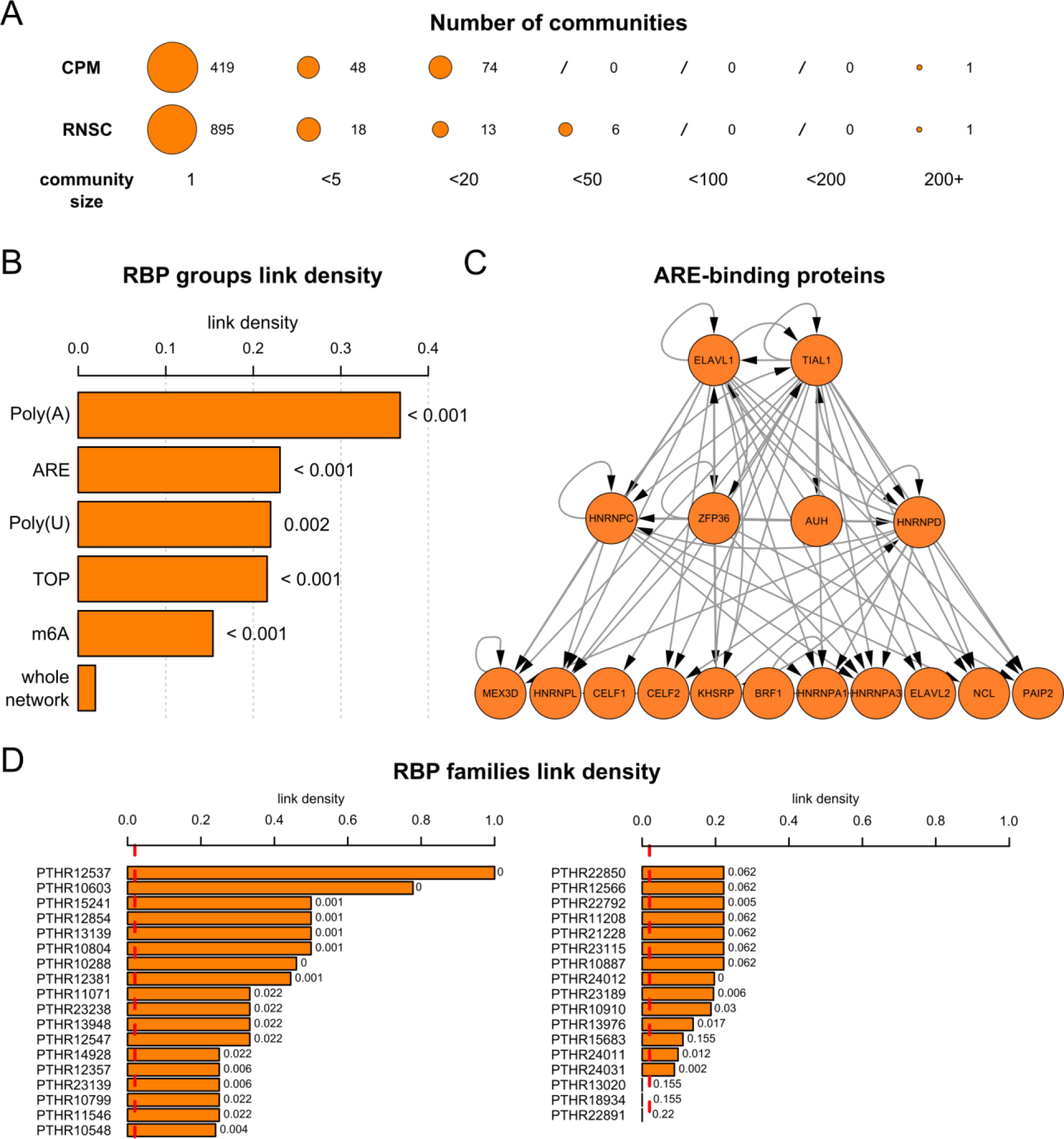
RBPs regulating common sets of targets are significantly more interconnected. **A)** displays the low modularity of the network as per the CPM and RNSC communities. Most are singletons, and one contains more than 25% of all RBPs. **B)** shows link density for the whole network and several groups of target-sharing RBPs: ARE, m6A, Poly(A), Poly(U) and TOP binding proteins; 1000-samples bootstrap p-values are shown next to each bar. **C)** shows the complete network of ARE-binding proteins, revealing a hierarchical structure dominated by *ELAVL1* and *TIAL1*. **D)** shows link density for families of RNA-binding proteins found in the network. A red dotted line indicates whole-network link density, and 1000-samples bootstrap p-values are shown next to each bar.

### RBP-RBP interactions occur in clusters dictated by their common target mRNAs

The number and size of the detected communities indicate a low modularity of the RBP-RBP network, likely due to a peculiar community structure which cannot be detected by current algorithms. To further study this aspect, we set out to investigate a more general principle, that of interactions between RBPs in the network being connected to cooperatively or competitively sharing mRNA targets. RBP-RBP network wiring constraints could indeed be due to combinatorial RBP interactions through their targets (both RBPs, which are in the network, and non-RBPs, which are outside it). We thus extracted all mRNA targets for each RBP in the network from the AURA 2 database (Dassi et al. 2014) and computed the overlap for every RBP pair. We compared these overlaps for protein-mRNA pairs in the network (interacting RBPs) and pairs not in the network (non-interacting RBPs). The results indicated that interacting RBPs share significantly more targets than non-interacting RBPs (median 141 and 52 resp., Wilcoxon test p<2.2E-16). To investigate the biological meaning of this general phenomenon, we then studied sets of RBPs known to bind to the same cis-element and consequently sharing most, if not all, of their targets. We considered AU-Rich Element (ARE) binding proteins (Barreau et al. 2005), proteins interacting with the 5’UTR terminal oligopyrimidine tract (TOP) element (Tcherkezian et al. 2014; Hamilton et al. 2006), proteins interacting with the m6A RNA modification sites ((Roignant and Soller 2017)), and finally proteins interacting with with poly(U) RNAs, and with the poly(A), a major cis-determinant of mRNA stability and translation (Goss and Kleiman 2013). ARE-binding proteins, in particular, are known to display both cooperative and competitive behaviors (Barreau et al. 2005). We computed link density (i.e. the fraction of all possible RBP-RBP interactions made within a group) for the whole network and each group. As shown in **Figure 2B**, all groups have significantly higher link densities than the whole network (7.8-18.7 times higher, 1000-samples bootstrap p-values=0.002 or less). The group with most interactions is the ARE-binding proteins (68 interactions), whose complete network is shown in **Figure 2C**. A hierarchical structure is visible, where *HuR*/*ELAVL1* and *TIAL1* are the major regulators (highest out-degree and lowest in-degree), connected to a second level (*ZFP36, HNRNPC*, *HNRNPD*, and *AUH),* which then controls the remaining RBPs (lowest out-degree).

Expanding on this idea, we eventually analyzed the link density of all annotated RBP families (as defined by Ensembl (Zerbino et al. 2018), see Methods). We assumed that, most often, members of the same RBP family cooperate or compete to regulate their common mRNA targets (Dassi 2017). Of the 288 families, 35 have at least two members in the network (i.e., taking part in at least one interaction). The median link density of these families is 0.24, with 32/35 having a higher density than the whole network (of these, 25 are significant according to a 1000-samples bootstrap) (**Figure 2D**). Although including only a fraction of all families, this results further indicate that RBP-RBP interactions may be needed to regulate cooperative and competitive behaviors on mutual targets, and that this behavior could be more prevalent than is currently known. “*RBP clusters*” (including families and sets of RBPs binding to the same cis-element) thus represent the community structure of the RBP-RBP network.

### RBP chains are master regulatory units of the cell

The interactions identified by analyzing RBP clusters are, however, only a fraction of all links in the network. We thus hypothesized that, alongside these community-like structures, the network could also be employing linear node chains as its functional units. To study this aspect, we extracted chains of length 4 and 5 (longest network path) from the network (examples are shown in **Figure 3A**). To assess their relevance, we checked whether chains were more functionally homogeneous (i.e. composed of RBPs with more closely related functions) than algorithm-derived communities, taken as comparison given their poor ability to capture the structure of the RBP-RBP network. We thus computed a *functional coherence* score as the average semantic similarity score for each pair of RBPs in a chain or community. Chains display a significantly higher functional coherence than algorithm-derived communities (Wilcoxon test p=9.01E-07/0.0347 for CPM/RNSC for chains of length 4; p=7.562E-06/0.086 for CPM/RNSC for length 5; shown as density in **Figure S2**). Chains thus seem to be relevant to the RBP-RBP network organization. In the TF-TF network, chains are instead significantly less coherent (average 0.41/0.43 vs. 0.75/0.73 for TF-TF and RBP-RBP of length 4 and 5 resp.; Wilcoxon test p<2.2E-16 for both lengths), suggesting a lesser importance of such units in this network.

**Figure 3:**
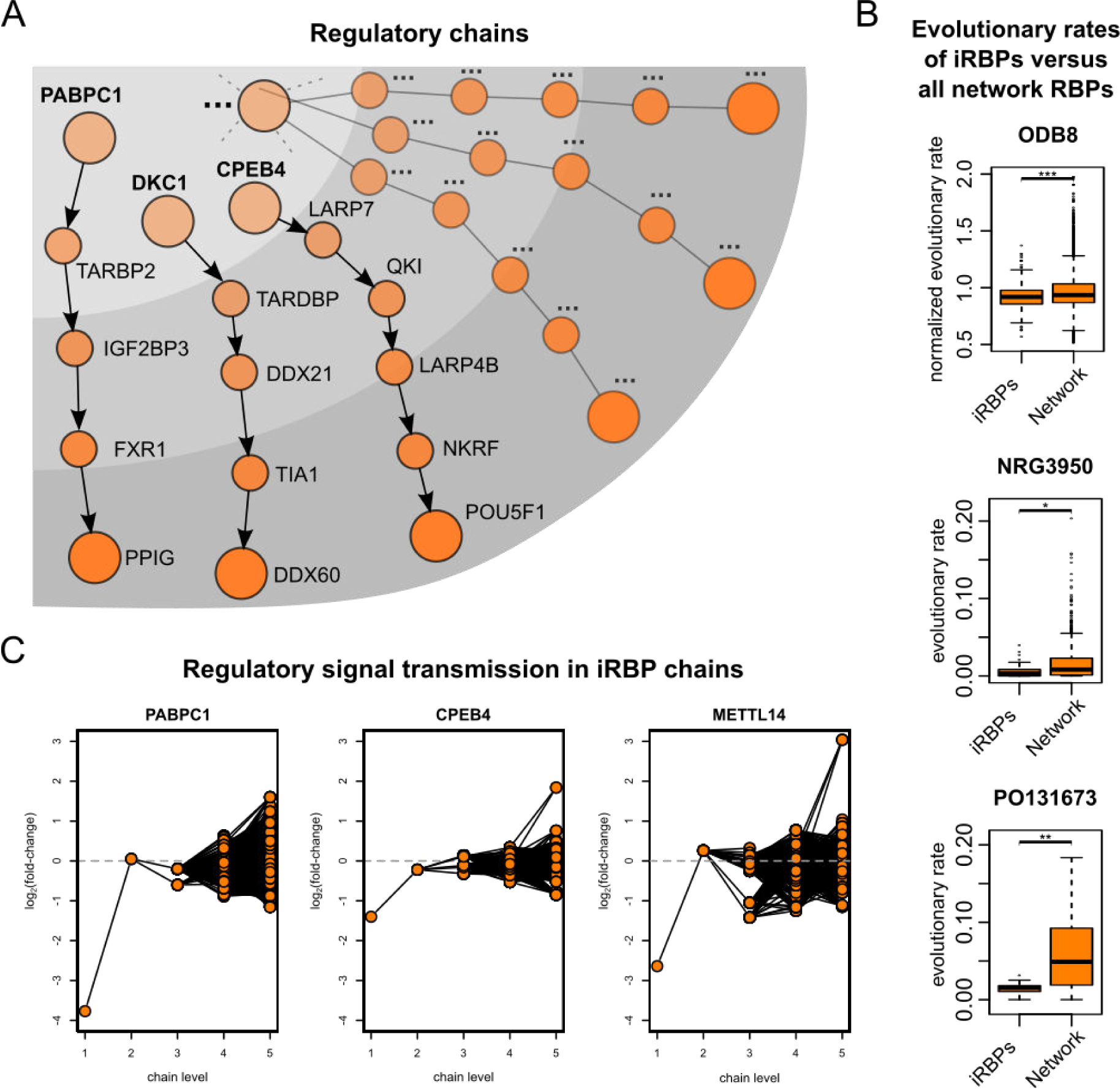
RBP chains dispatch regulatory information in the network. **A)** examples of RBP chains. Dashed lines and dotted name represent an iRBP heading many RBP chains. Increasing node color intensity represents the transmission of regulatory input through the chain, from the first to the last node. **B)** evolutionary rates of iRBPs and all RBPs in the network, obtained from the ODB8 database and two articles. iRBPs have a significantly lower rate in all datasets (Wilcoxon test p=5.2E-05, 0.0128 and 0.0016 for ODB8, NRG3950, and PO131673). **C)** Displays the log2 fold-change for RBPs at the various levels of chains led by PABPC1, CPEB4, and METTL14 when silencing these iRBPs. The first level of the chain is the silenced iRBPs, while level 5 represents the last RBP of a chain (with levels 2..4 being intermediate steps of each chain). Lines represent the RBP-RBP connections in a chain while orange circles represent RBPs.

Chains are headed by a few initiator RBPs (iRBPs, 53 genes): these could be the most influential regulators, able to control many other RBPs and processes to dictate cell phenotypes. Therefore, iRBPs could be essential for the proper execution of cell processes. On this assumption, we searched for iRBPs in essential genes (defined by the underrepresentation of gene-trap vectors integration in their locus) of two human cell lines, as per a recent work (Blomen et al. 2015). As shown in **Table S1**, a third of the iRBPs are essential in both cell lines, with 21 (40%) essential in the HAP1 line. iRBPs are enriched for essential genes in these cell models (max. Fisher test p=1.73E-05), and a 1000-samples bootstrap was significant (p<0.001) in both cell lines and their intersection. Most iRBPs (43/53, 81%) are also essential in at least one cellular model as per RNAi screenings included in the GenomeRNAi database (Schmidt et al. 2013). Merging all these annotations yields the remarkable total number of 46/53 iRBPs essential in at least one cell model (86%). To further strengthen this finding, we obtained the orthologs of iRBPs in mouse, D. *melanogaster,* and C. *elegans*, and compared them with essential genes in those organisms. As shown in **Table S5**, **S6**, and **S7**, the enrichment of essential genes in iRBPs is highly significant also for these organisms.

The iRBPs could also be highly conserved, due to their fundamental role in driving the RBP chains. We thus investigated whether these RBPs are more evolutionarily constrained than other RBPs. We extracted evolutionary rates of sequence divergence from the ODB8 database (Zdobnov et al. 2017) and (Zhang and Yang 2015), and rates of purifying selection from (Kryuchkova-Mostacci and Robinson-Rechavi 2015). We observed that, in all datasets, iRBPs have a significantly lower evolutionary rate than all RBPs in the network (**Fig 3B**; Wilcoxon test p=5.2E-05, 0.0128 and 0.0016 for ODB8, NRG3950 and PO131673 respectively). Furthermore, we also investigated whether the iRBPs PTR is evolutionarily constrained. We thus computed their average UTR conservation, and first found that UTRs of RBPs in the network are more conserved than the UTRs of all genes (Wilcoxon test p < 2.2E-16 for 5’ and 3’ UTRs). As the network includes most RBPs, this feature is characteristic of RBP genes. iRBPs 3’UTR conservation was also found to be significantly higher than that of other RBPs (Wilcoxon test p=0.002142). This ultra-conservation, coupled with the essentiality of most iRBPs, consistently support their importance as key cell regulators.

We eventually asked ourselves whether the regulatory information is transmitted through the chains, from the iRBPs down to the last node. To study this aspect, we obtained and reanalyzed publicly available transcriptome profiles of knock-down experiments for three iRBPs (two with chains of length 4 and 5, *PABPC1* and *CPEB4*, and one with chains of length 4, *METTL14,* see Methods) in human cells. These RBPs act on various processes, including, for PABPC1 and METTL14, the regulation of mRNA stability. (Weng et al. 2018; Wang and Kiledjian 2000). We thus expect to detect at least a partial effect of their knock-down on these transcriptomes. We plotted the fold-change (knock-down versus control) of the RBPs composing the chains controlled by each of the three iRBPs. As shown in **Figure 3C**, a sizeable fraction of all chain members are differentially expressed (23.9% for PABPC1, 22.4% for CPEB4, and 46.3% for METTL14 at the adjusted p-value threshold of 0.05); if considering only a permissive fold-change threshold of 1.1 these numbers rise two to three times (66.1% for PABPC1, 44.1% for CPEB4, and 61.9% for METTL14). It must be noted that other modes of regulation, which cannot be observed in these datasets, can also be used by these proteins aside from mRNA stability (e.g., translational control). This data thus suggest that the regulatory information sparked by an iRBP is indeed transmitted through its chains, likely expanding the set of processes which can be controlled by these proteins. Chains are thus a functional unit in the RBP-RBP network, complementing the observed RBP clusters.

### The RBP-RBP network is a robust and efficient hierarchy

We finally asked ourselves which were the implications of RBP chains on the global network structure. A reasonable hypothesis is that chains induce a hierarchical structure, as also suggested by the ranked clusters model we observed as defining the local network structure. We thus measured how hierarchical is the RBP-RBP network (Cheng et al. 2015), which revealed it as much more than any of the 41 TF-TF networks. When considering a hierarchy of 2, 4 or 6 levels; p-value is always orders of magnitude lower, with a −log_10_p of 14.2 versus an average of 3.85 for TF-TF networks at six levels. Furthermore, feedback loops (not coherent with a hierarchical organization) are depleted in the network, representing 0.0085% of the motifs only; feed-forward loops, coherent with a hierarchical organization, are instead enriched and amount to 3.29% of the motifs.

We then assessed another desirable property, that of network robustness to the “removal” (i.e., loss of function) of an RBP from the network. To do so, we computed the pairwise disconnectivity metrics on each node (Potapov et al. 2008). The metric is low (only 0.14% of pairs are disconnected on average when removing a node from the network) and significantly lower than for the TF-TF networks (average is three times higher for TFs, worst p-value=5.6E-104). The network is thus well-tolerant to losing a node (fewer nodes are disconnected when removing a node), which implies that RBP-RBP interactions are robust. This feature is likely granted by the use of densely connected RBP clusters, resulting in partially redundant regulation.

Eventually, while RBP clusters are redundant by definition (as they co-regulate a largely overlapping set of targets), we asked whether also single RBP chains shared this property. We thus computed the overlap between all targets (both RBPs, which are in the network, and non-RBPs, which are outside it) of RBPs at the various levels of each chain of length 5. It resulted being particularly low, as only 7.6% of the targets are overlapping between any two levels (median of all chains, average of each pair in a chain; the range is 2.8%-15.5%). Differently from RBP clusters, we can thus say that chains are efficient, as targets are not redundantly regulated by individual RBPs along the chain, but rather are predominantly organized in complementary sets at each of its nodes. This efficiency comes at the expense of robustness (i.e., if one level of the chain fails the regulatory signal would most often be lost), which is instead a feature of RBP clusters. The resulting model, shown in **Figure 4**, couples hierarchical structure, network robustness through RBP groups, and efficiency through RBP chains.

**Figure 4:**
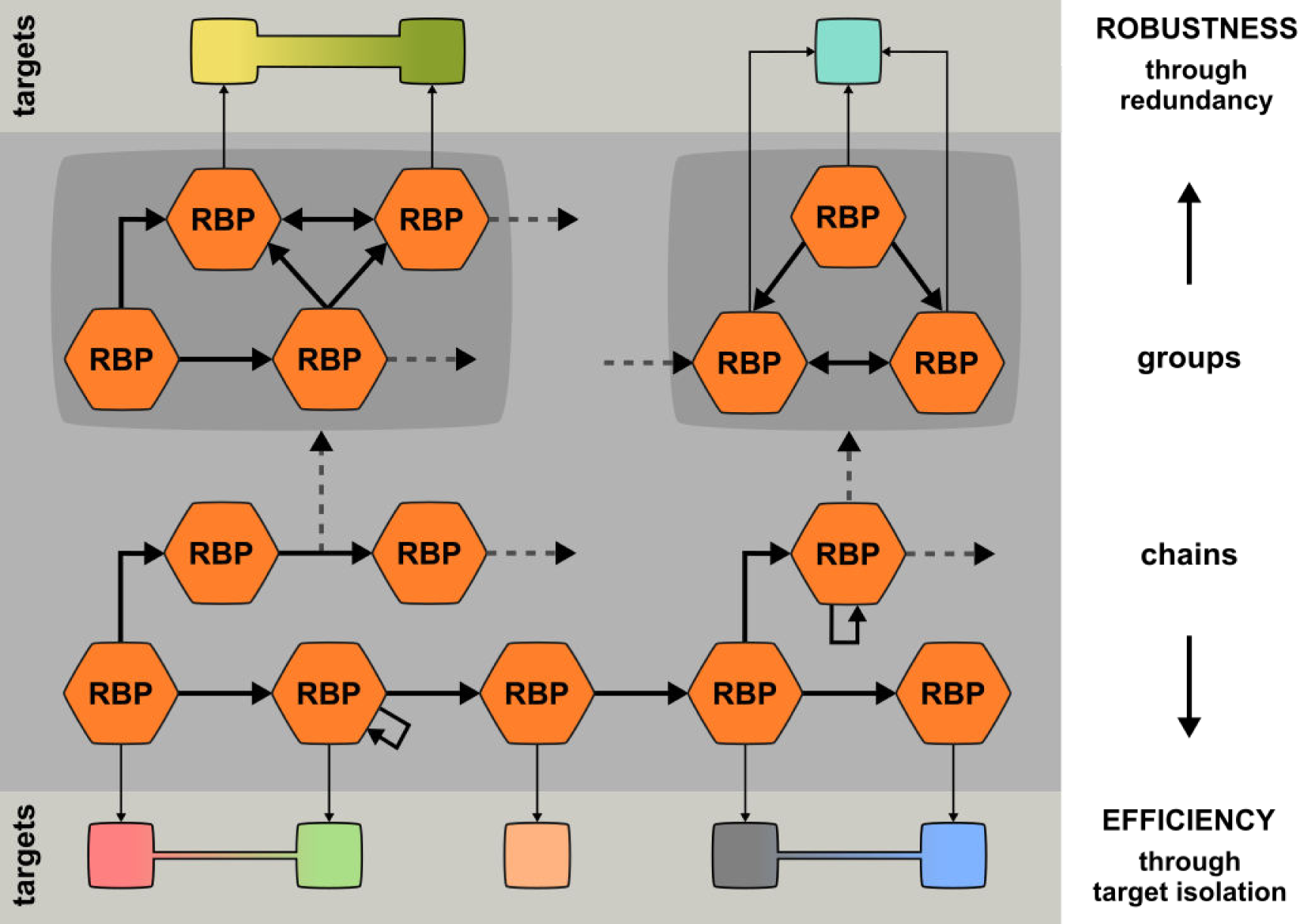
The RBP-RBP network is a robust and efficient hierarchy. Model of the RBP-RBP network as derived from our analyses. RBPs are indicated by hexagon-shaped nodes, RB-RBP interactions by thick arrows (arrows pointing to the originating RBP represent autoregulation events), and targets sets by squares which can be shared by multiple RBPs (fraction of shared targets represented by the size of the shared area between the two squares). RBP-RBP interactions are robust due to densely connected RBP groups (co-regulating most of their targets), while RBP chains confer hierarchy and efficiency to the network, as target mRNA sets for each RBP in a chain are completely or predominantly different (“isolated”). Dashed in and out-going arrows hint to the presence of further interactions within and between RBP groups and chains in the network.

## Discussion

We presented here the first characterization of the RBP-RBP regulatory network. Starting from several reports hinting to a post-transcriptional hierarchy of regulators (Potapov et al. 2008; Dassi et al. 2013; Mukherjee et al. 2011; Pullmann et al. 2007), we collected available RBP-mRNA association data and described the network of interactions involving an RBP and an RBP mRNA. The network is small-world and scale-free, typical properties of gene regulatory networks. While the network is partial (as data is available for a fraction of all RBPs only), its local motif structure is highly coherent with the one of the inferred network, suggesting that it is representative of general patterns in RBP-RBP interactions.

Its local structure is similar to the one of TF-TF networks derived by DNase footprinting (Neph et al. 2012). However, differential enrichment of several motifs suggests that structure specialization occurs in the RBP-RBP network with respect to the TF-TF one, likely aimed at better suiting the specificities of post-transcriptional regulation. In particular, we found up- and down-linked mutual dyads as particularly enriched. These motifs are distinctive of a ranked clusters structure (Johnsen 1985), thus suggesting that the network can be divided in features conferring hierarchy and clusters of densely interacting nodes.

To study the role of these interactions in shaping cell phenotypes, we investigated why RBPs regulate each other. We found a few protein complexes involved in RNA metabolism and highly intra-regulated by RBP-RBP interactions. However, only a fraction of all complexes display this behavior, which cannot thus be considered general. We instead observed that groups of RBPs having overlapping targets tend to regulate each other: these interactions could represent a novel layer of regulation for cooperative and competitive behaviors between RBPs. As known for ARE-binding proteins (Barreau et al. 2005), RBPs can tune the expression of a common target by competitive or cooperative binding. We suggest that RBPs may influence the outcome of this process also by regulating the expression of the partner RBP, a mechanism which could be used to reach precise ratios in a mRNP and yield the intended regulatory effect on common targets.

These groups represent “islands” of densely connected RBPs and are key in providing robustness to the network. Indeed, their partially redundant regulation improves the resilience of the network to the loss of function of individual RBPs.

However, RBP groups are not the only constituent feature of this network. Studying how RBPs interact with each other, we uncovered a set of widespread linear structures which appear to be more prevalent than communities. These structures, which we termed RBP chains and are driven by a few initiator RBPs (iRBPs), could provide enhanced flexibility with respect to a community pattern. Indeed, we believe that RBPs evolved the ability to influence a broad set of biological processes through such chains. Most iRBPs are essential for the cell, their 3’UTRs are more conserved, and their evolutionary rates are lower than for other RBPs. Taken together, these findings truly back iRBP importance as ancient master regulators of cell processes.

Chains profoundly shape the RBP-RBP network to be highly hierarchical. Their regulatory action confers efficiency, as the fraction of targets shared between the different chain levels is limited (i.e., regulation is not replicated along a chain), thus streamlining the flow of regulatory information from iRBPs to the final chain targets.

We have thus identified the two features hypothesized by the ranked clusters model: a hierarchy-inducing structure (the RBP chains), and clusters of densely interacting nodes (RBP groups). This indicates that this model fits well with the RBP-RBP network and can be found at different depths of observation: from the local, three-nodes motif structure, to the patterns defining the topology of the global network. The combination of properties offered by these features, namely robustness and efficiency, reflects the constant evolutionary pressure shaping a machinery as fundamental to the cell as is the one driving post-transcriptional regulation of gene expression. While establishing robustness only through RBP groups could in principle lead to a weaker architecture (as the regulatory signal going through chains is not replicated), it may be cheaper to obtain and potentially more far-reaching. This consideration, however, raises a question: which is the role of chains in relation to RBP groups?

We suggest that RBP chains use the modulation of RBP targets as a connector to different processes, represented by the RBP groups. We call this model “*island-hopping*”. Under this model, the regulatory signal originated from the chains iRBPs hops from one island (RBP group, side-connected to the chain) to another, while flowing through the chain levels, to regulate several cellular processes. This pattern thus allows potentially coordinating a broad set of functions of interest. Activating different chains would then result in the modulation of a different set of processes, granting substantial flexibility to the RBP-RBP network. This work thus establishes interactions among RBPs and RBP mRNAs as a backbone driving post-transcriptional regulation of gene expression to coordinately tune protein abundances.

## Materials and Methods

### RBP list construction and annotation

We built the list of human RNA-binding proteins by first extracting genes annotated as RNA-binding (GO:0003723) and protein-coding in Ensembl v92 (Zerbino et al. 2018), then merging these genes with the curated RBP list from a recent work (Sebestyén et al. 2016). The resulting list thus includes canonical and novel RBPs for a total of 1827 proteins. Families of RNA-binding proteins were extracted from Ensembl v92 gene families (Zerbino et al. 2018), by considering only families including more than one RBP.

### Network construction

Regulatory interactions involving two RBPs were extracted from the AURA 2 database (Dassi et al. 2014). Interactions were filtered by requiring the expression of both participants in HEK293, HeLa or MCF7 cells, systems where the majority of the data were derived. Expressed genes were determined by RNA-seq profiles of HEK293 (Kishore et al. 2011), HeLa (Cabili et al. 2011) and MCF7 (Vanderkraats et al. 2013), using an expression threshold of 0.1 RPKM. The direction of edges in the network represents regulation by the source RBP on the target RBP mRNA.

The inferred RBP-RBP network was built by collecting RBP-bound regions in mRNA UTRs from a protein occupancy profiling assay in HEK293 cells (Dassi et al. 2014; Baltz et al. 2012). RNAcompete-derived PWMs for 193 human RBPs (Ray et al. 2013) were downloaded from CISBP-RNA (Ray et al. 2013). Binding regions of these RBP on protein-bound regions were identified by Biopython (Ray et al. 2013; Cock et al. 2009), selecting the best matching RBP for each region (score threshold=0.99); only interactions involving two RBPs (one binding to the other mRNA) were included in the network.

The networks were deposited in NDEX with ID *ee3e8898-6e29-11e8-a4bf-0ac135e8bacf*, *f5ad750b-6e29-11e8-a4bf-0ac135e8bacf* and *fc1e526e-6e29-11e8-a4bf-0ac135e8bacf*.

### Network properties analysis

Network diameter, degree distribution, closeness centrality, Watts-Strogatz (CC1) and two-neighbor (CC2) clustering coefficient were computed by Pajek (Batagelj and Mrvar 2002) and plotted with R (Tierney 2012). The S^WS^ measure was computed as described in (Humphries and Gurney 2008) by using the Watts-Strogatz clustering coefficient and generating the required random network with Pajek (Batagelj and Mrvar 2002). The network control structure was computed by Zen (Ruths and Ruths 2014). Hierarchical score was computed as per (Cheng et al. 2015) and pairwise disconnectivity obtained by DiVa (Potapov et al. 2008).

Link density for a set of nodes was computed as (number of links between nodes in the set) / (number of nodes in the set^2). Bootstraps were performed by 1000 random selections of a number of nodes equal to the set size and computation of the link density for each of these.

### Network structure analysis

Network motifs of size 3 and 4 were identified with FANMOD (Cheng et al. 2015; Wernicke and Rasche 2006) using 1000 random networks (100 for motifs of size 4, due to required computing time), 3 exchanges per edge and 3 exchange attempts. Triad significance profiles for motifs of size 3 were computed as described in (Milo et al. 2004) for the RBP-RBP network, the inferred RBP-RBP network and the TF-TF networks described in (Neph et al. 2012).

Communities were studied with the SurpriseMe tool (Aldecoa and Marin 2013): CPM (Palla et al. 2005) and RNSC (King et al. 2004), the algorithms obtaining the highest S values, were used to define communities. Chains were extracted from the network with igraph (http://igraph.org); functional coherence scores were computed with GOSimSem (Yu et al. 2010) by averaging the semantic similarity of each pair of genes in the chain.

### Protein-protein interactions and complexes overlap

Human protein-protein interactions were extracted from STRING (Szklarczyk et al. 2017), BioPlex (Huttlin et al. 2017), and IntAct (Orchard et al. 2014), retaining only interactions of the “binding” type (physical association) and with both partners being in our network. Human protein complexes were downloaded from CORUM (Ruepp et al. 2010). Overlaps were performed by custom Python scripts.

### Gene essentiality and phylogenetic conservation analysis

Essential genes of human cells were obtained from (Blomen et al. 2015). Genes associated with an embryonic lethal phenotype in mouse from the MGI (Blake et al. 2017); genes associated with a lethality phenotype were extracted from WormBase (Lee et al. 2018) and FlyBase (Gramates et al. 2017) for C.*elegans* and D.*melanogaster* respectively. Orthologs of iRBPs were extracted from the same databases. Bootstraps were computed by 1000 random selections of as many genes as iRBPs and computing the fractions of these in the essential genes for each organism.

UTR conservation scores were computed by averaging the phastCons score derived from the UCSC 46-way vertebrate alignment (Casper et al. 2018). The average score of all 5’ or 3’ UTRs of a gene was employed as the conservation score for that gene 5’ or 3’UTRs. Protein evolutionary rates were obtained from the ODB8 database (Zdobnov et al. 2017) and two articles (Kryuchkova-Mostacci and Robinson-Rechavi 2015; Zhang and Yang 2015); statistical tests were performed by R (Tierney 2012).

### iRBP knock-down datasets

RNA-seq datasets following the knock-down of PABPC1, CPEB4, and METTL14 were obtained from GEO (IDs: GSE88099, GSE88545, and GSE56010). Reads were quality-trimmed and adapters removed with Trimmomatic (Bolger et al. 2014), then aligned to the human genome (hg38 assembly), and transcripts quantified (Gencode v28 annotation) with STAR (Dobin et al. 2013). Differential expression was eventually computed with DESeq2 (Love et al. 2014) using an adjusted p-value threshold of 0.05.

## Supplemental Figures

**Figure S1:**
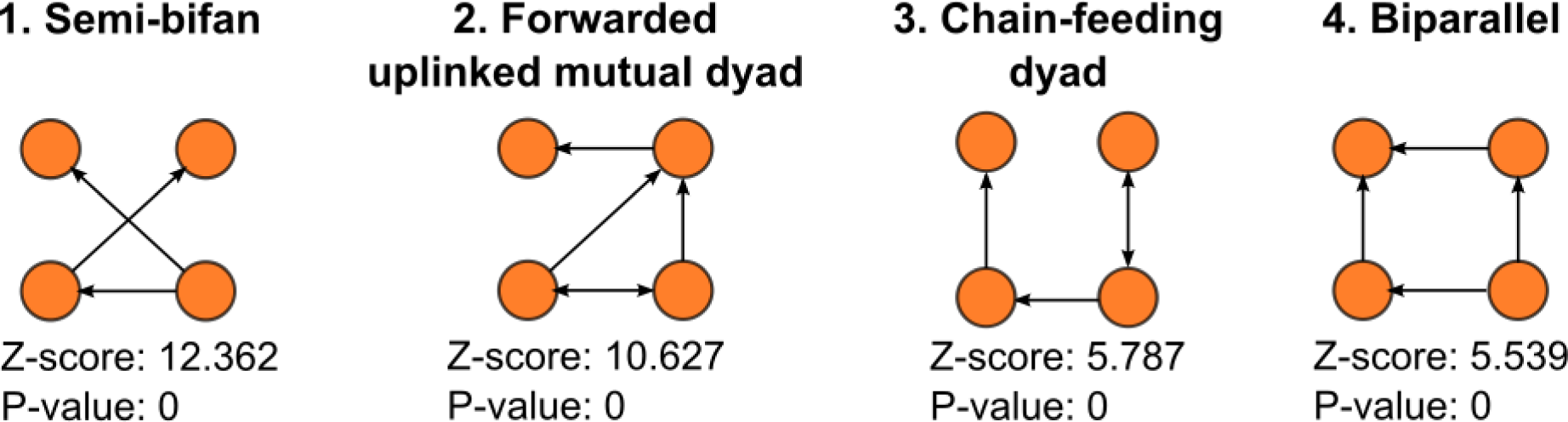
Most enriched 4-nodes motifs in the network. Shows the four most significant 4-nodes motifs identified by FANMOD with their z-score and p-value.

**Figure S2:**
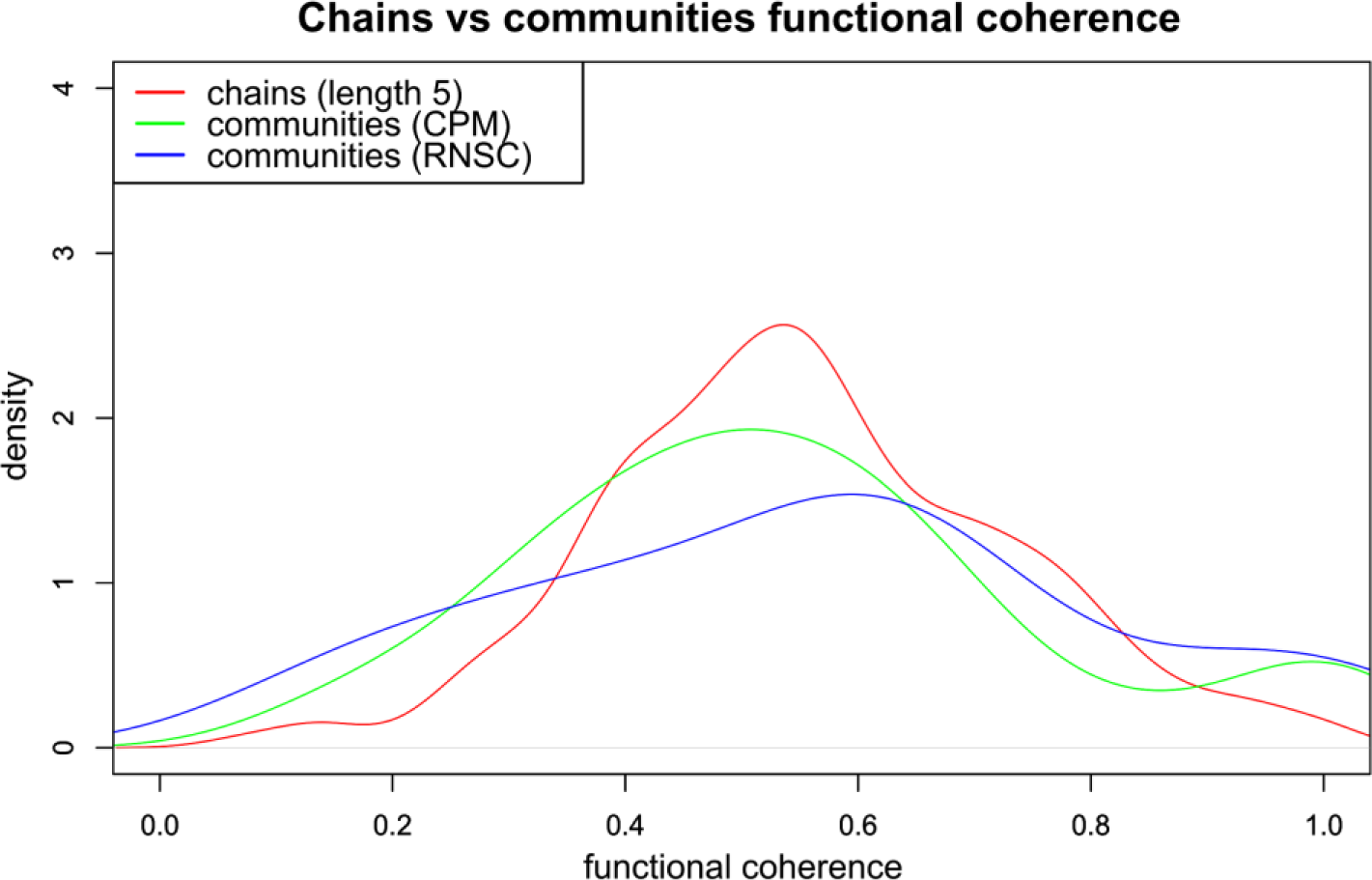
Functional coherence distribution for chains versus communities. Shows the differences in density of the functional coherence values distribution between chains of length 6 (in red) and communities obtained by CPM (green) and RNSC (blue).

